# 14-3-3 phosphorylation inhibits 14-3-3θ’s ability to regulate LRRK2 kinase activity

**DOI:** 10.1101/2023.05.27.542591

**Authors:** Rudradip Pattanayak, Chad M. Petit, Talene A. Yacoubian

## Abstract

LRRK2 mutations are among the most common genetic causes for Parkinson’s disease (PD), and toxicity is associated with increased kinase activity. 14-3-3 proteins are key interactors that regulate LRRK2 kinase activity. Phosphorylation of the 14-3-3θ isoform at S232 is dramatically increased in human PD brains. Here we investigate the impact of 14-3-3θ phosphorylation on its ability to regulate LRRK2 kinase activity. Both wildtype and the non-phosphorylatable S232A 14-3-3θ mutant reduced the kinase activity of wildtype and G2019S LRRK2, whereas the phosphomimetic S232D 14-3-3θ mutant had minimal effects on LRRK2 kinase activity, as determined by measuring autophosphorylation at S1292 and T1503 and Rab10 phosphorylation. However, wildtype and both 14-3-3θ mutants similarly reduced the kinase activity of the R1441G LRRK2 mutant. 14-3-3θ phosphorylation did not promote global dissociation with LRRK2, as determined by co-immunoprecipitation and proximal ligation assays. 14-3-3s interact with LRRK2 at several phosphorylated serine/threonine sites, including T2524 in the C-terminal helix, which can fold back to regulate the kinase domain. Interaction between 14-3-3θ and phosphorylated T2524 LRRK2 was important for 14-3-3θ’s ability to regulate kinase activity, as wildtype and S232A 14-3-3θ failed to reduce the kinase activity of G2019S/T2524A LRRK2. Molecular modeling showed that 14-3-3θ phosphorylation causes a partial rearrangement of its canonical binding pocket, thus affecting the interaction between 14-3-3θ and the C-terminus of LRRK2. We conclude that 14-3-3θ phosphorylation destabilizes the interaction of 14-3-3θ with LRRK2 at T2524, which consequently promotes LRRK2 kinase activity.

## INTRODUCTION

Mutations in Leucine-rich repeat kinase 2 (LRRK2) are recognized among the most common genetic causes of Parkinson disease (PD) [1–3]. LRRK2 is a large multi-domain protein which includes an ankyrin repeat (ANK), an armadillo repeat (ARM), a leucine rich repeat (LRR) domain, a kinase domain, a RAS domain, a GTPase domain, and a WD40 domain [4]. Several mutations reported for LRRK2 have been found to be pathogenic: R1441G/C/H within the ROC domain; Y1699C within the COR domain; and I2012T, N1437H, G2019S, and I2020T within the kinase domain [5]. All prominent mutations localize in the enzymatic core of LRRK2, spanning through the GTPase and kinase domains, and can alter LRRK2 enzymatic activity [4, 6–9]. Among these, G2019S is the most common LRRK2 mutation associated with PD and is associated with increased LRRK2 kinase activity [1, 6–8, 10, 11].

Key interactors for LRRK2 are the highly conserved family of 14-3-3 proteins [12–14]. 14-3-3s are highly-expressed brain chaperone proteins that interact with a diverse group of signaling proteins including kinases [15]. There are seven 14-3-3 isoforms (14-3-3 β, γ, ε, η, ζ, σ, and τ/θ), and the majority of 14-3-3 isoforms have been shown to interact with LRRK2 [12, 14]. 14-3-3s dimerize to form amphipathic binding sites to interact with a wide host of ligands, including LRRK2, and many protein-protein interactions with 14-3-3s are regulated by phosphorylation of the ligand [15, 16]. We previously reported that the 14-3-3θ isoform is a major regulator for LRRK2’s kinase activity [17]. 14-3-3θ also prevents neurite shortening induced by either the G2019S or R1441G LRRK2 mutant [17]. The regulatory effect of 14-3-3θ on LRRK2 kinase activity is dependent on direct interaction between LRRK2 and 14-3-3θ, as disruption of the interaction prevented such regulation [17]. Several different LRRK2 sites are implicated in binding to 14-3-3s, including S910, S935, S1444, and T2524 [12, 14, 18, 19].

Alterations in 14-3-3 proteins have been observed in human PD. Several 14-3-3 isoforms, including 14-3-3θ, are found within Lewy Bodies, the key pathological hallmark of PD [20–22]. We also recently reported a dramatic increase in 14-3-3θ phosphorylation at S232 in detergent insoluble fractions from human PD and Dementia with Lewy bodies (DLB) brains, and this phosphorylation correlates with cognitive decline and pathological severity [23]. 14-3-3θ phosphorylation at S232 inhibits the neuroprotective effect of 14-3-3θ against mitochondrial toxins associated with increased PD risk [24]. Here we hypothesize that increased 14-3-3θ phosphorylation at S232 may disrupt its ability to regulate LRRK2 kinase activity.

In this study, we investigated the effect of S232 phosphorylation of 14-3-3θ on wildtype (WT) and two different LRRK2 mutants, G2019S and R1441G. We found that 14-3-3θ phosphorylation disrupted 14-3-3θ’s ability to regulate both WT and G2019S LRRK2 kinase activity, but not R1441G mutant. While 14-3-3θ phosphorylation did not disrupt global binding to LRRK2 as determined by immunoprecipitation and proximity ligation assays (PLA), we observed that S232 phosphorylation promoted conformational alterations in the binding interaction with LRRK2 involving T2524 at the C-terminal end in order to disrupt kinase regulation.

## RESULTS

### 14-3-3θ phosphorylation at S232 reduces the ability of 14-3-3θ to reduce LRRK2 autophosphorylation

We have previously shown that the 14-3-3θ isoform can reduce WT and mutant LRRK2 kinase activity as measured by autophosphorylation at S1292 and T1503 [17]. To monitor the effect of 14-3-3θ phosphorylation on LRRK2 kinase activity, we tested the impact of the non-phosphorylatable S232A 14-3-3θ mutant and the phosphomimetic S232D 14-3-3θ mutant on WT and mutant LRRK2 autophosphorylation. We co-transfected FLAG-tagged G2019S LRRK2 with either empty vector (EV), V5-tagged WT 14-3-3θ, or V5-tagged S232 14-3-3θ mutants in HEK293T cells and then measured LRRK2 autophosphorylation at S1292 (p-S1292). As observed previously, WT 14-3-3θ significantly decreased S1292 phosphorylation of G2019S LRRK2 by 28% (Fig. 1A). The S232A mutant was more effective at reducing S1292 phosphorylation of G2019S LRRK2 by 42%, but S232D was less effective at reducing S1292 phosphorylation of G2019S LRRK2 (Fig. 1A).

**Figure 1.**
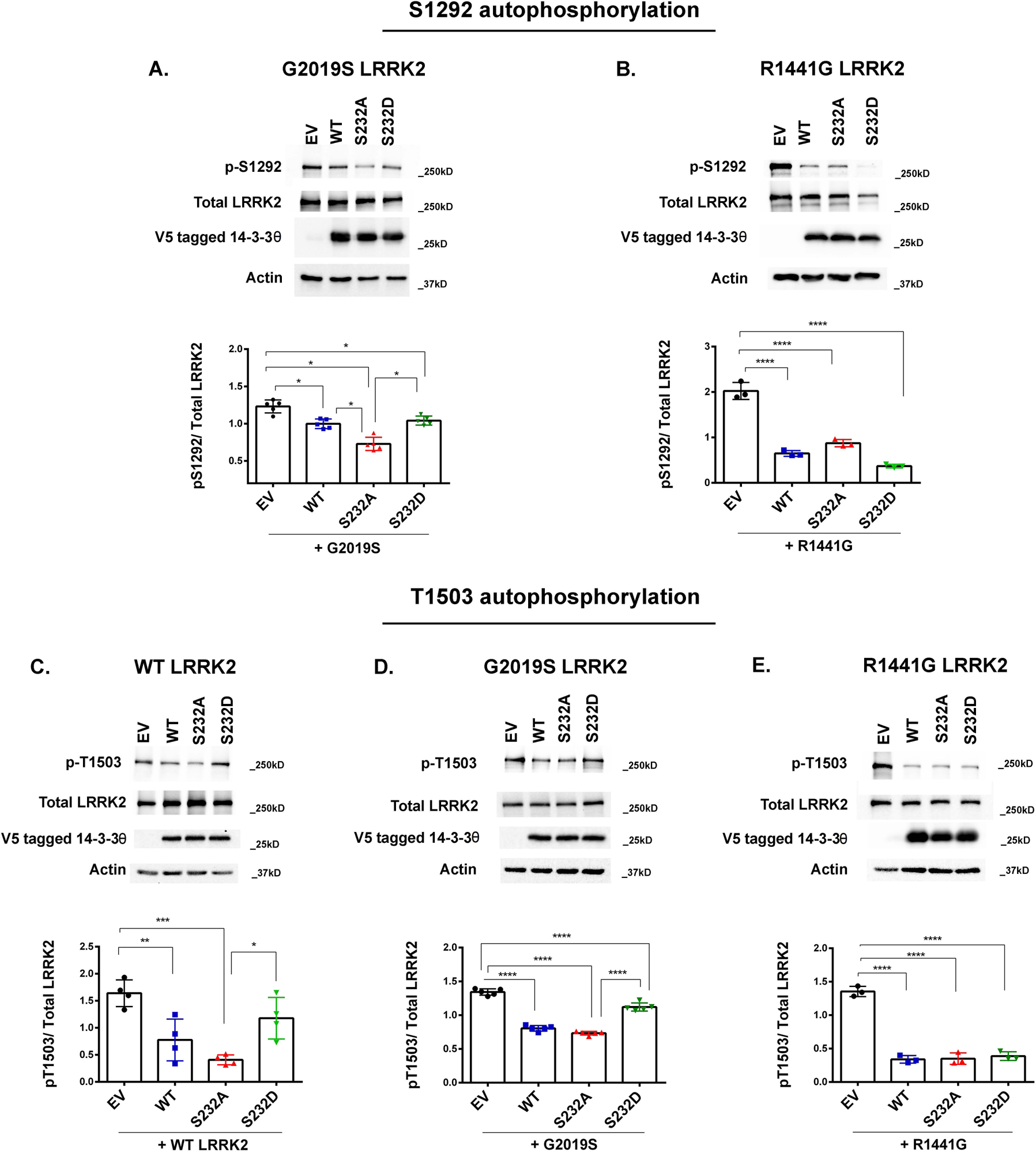
S232 phosphorylation disrupts the ability of 14-3-3θ to reduce autophosphorylation of WT and G2019S LRRK2. **(A)** Representative Western blot and quantification of p-S1292 LRRK2 levels in lysates of HEK 293T cells co-transfected with empty vector (EV), WT, S232A, or S232D 14-3-3θ with G2019S LRRK2. n = 5 independent rounds. *p < 0.05 (Tukey’s multiple comparison test). **(B)** Representative Western blot and quantification of p-S1292 LRRK2 levels in lysates of HEK 293T cells co-transfected with EV, WT, S232A, or S232D 14-3-3θ with R1441G LRRK2. n = 3 independent rounds. **** p < 0.0001 (Tukey’s multiple comparison test). **(C-E)** HA-tagged WT (C), HA-tagged G2019S (D), and c-myc-tagged R1441G LRRK2 (E) were co-transfected with EV, WT, S232A or S232D 14-3-3θ into HEK 293T cells, and then LRRK2 was immunoprecipitated with respective HA and c-myc antibodies to isolate LRRK2 for kinase reactions. Following kinase reaction, immunoprecipitants were analyzed for p-T1503 and total LRRK2 by Western blot. n = 4 independent rounds for WT LRRK2, n = 5 independent rounds for G2019S, and n = 3 independent rounds for R1441G. *p < 0.05, **p < 0.01, ***p < 0.001, ****p < 0.0001 (Tukey’s multiple comparison test).

We next tested the impact of S232 mutants to reduce the kinase activity of LRRK2 with the R1441G mutation, which is found in the GTPase domain instead of the kinase domain. We transfected myc-tagged R1441G with EV, WT or S232 14-3-3θ mutants and measured S1292 LRRK2 phosphorylation. We found that WT, S232A, and S232D 14-3-3θ all reduced S1292 phosphorylation of R1441G to a similar degree (Fig. 1B).

To validate our findings above, we next tested the impact of S232 phosphorylation on LRRK2 kinase activity using a different LRRK2 kinase assay involving the T1503 autophosphorylation site (Fig. 1C-E). In this assay, we performed an *in vitro* kinase assay after pulldown for LRRK2 from transfected HEK 293T cells and measured p-T1503 levels.

We found that WT and S232A 14-3-3θ both significantly reduced p-T1503 levels of WT LRRK2, while the S232D mutant did not significantly reduce p-T1503 levels (Fig. 1C). Similar effects were seen with G2019S LRRK2: WT and S232A 14-3-3θ mutant significantly reduced p-T1503 levels of G2019S, while S232D 14-3-3θ showed a slight reduction in p-T1503 levels (Fig. 1D). In the case of R1441G LRRK2, WT, S232A, and S232D 14-3-3θ reduced T1503 phosphorylation equally (Fig.1E), similar to the findings for p-S1292 (Fig. 1B).

### 14-3-3θ phosphorylation’s effect on LRRK2-mediated Rab10 phosphorylation

We next examined the impact of 14-3-3θ phosphorylation on the kinase activity of LRRK2 towards other kinase substrates besides itself. LRRK2 phosphorylates several Rab proteins, including Rab8, Rab10 and Rab12 [25–27]. We tested the ability of WT and mutant 14-3-3θ to reduce Rab10 phosphorylation induced by G2019S LRRK2. HEK293T cells were transfected with HA-tagged Rab10, G2019S LRRK2, and with either EV, WT 14-3-3θ, or mutant S232 14-3-3θ. We found that WT 14-3-3θ significantly reduced Rab10 phosphorylation by 39% (Fig. 2). The S232A mutant caused a 67% reduction in Rab10 phosphorylation, but S232D 14-3-3θ failed to reduce Rab10 phosphorylation induced by G2019S (Fig. 2). Based on these findings, we concluded that 14-3-3θ phosphorylation at S232 prevents 14-3-3θ’s regulation of LRRK2 kinase activity towards itself and Rab substrates.

**Figure 2.**
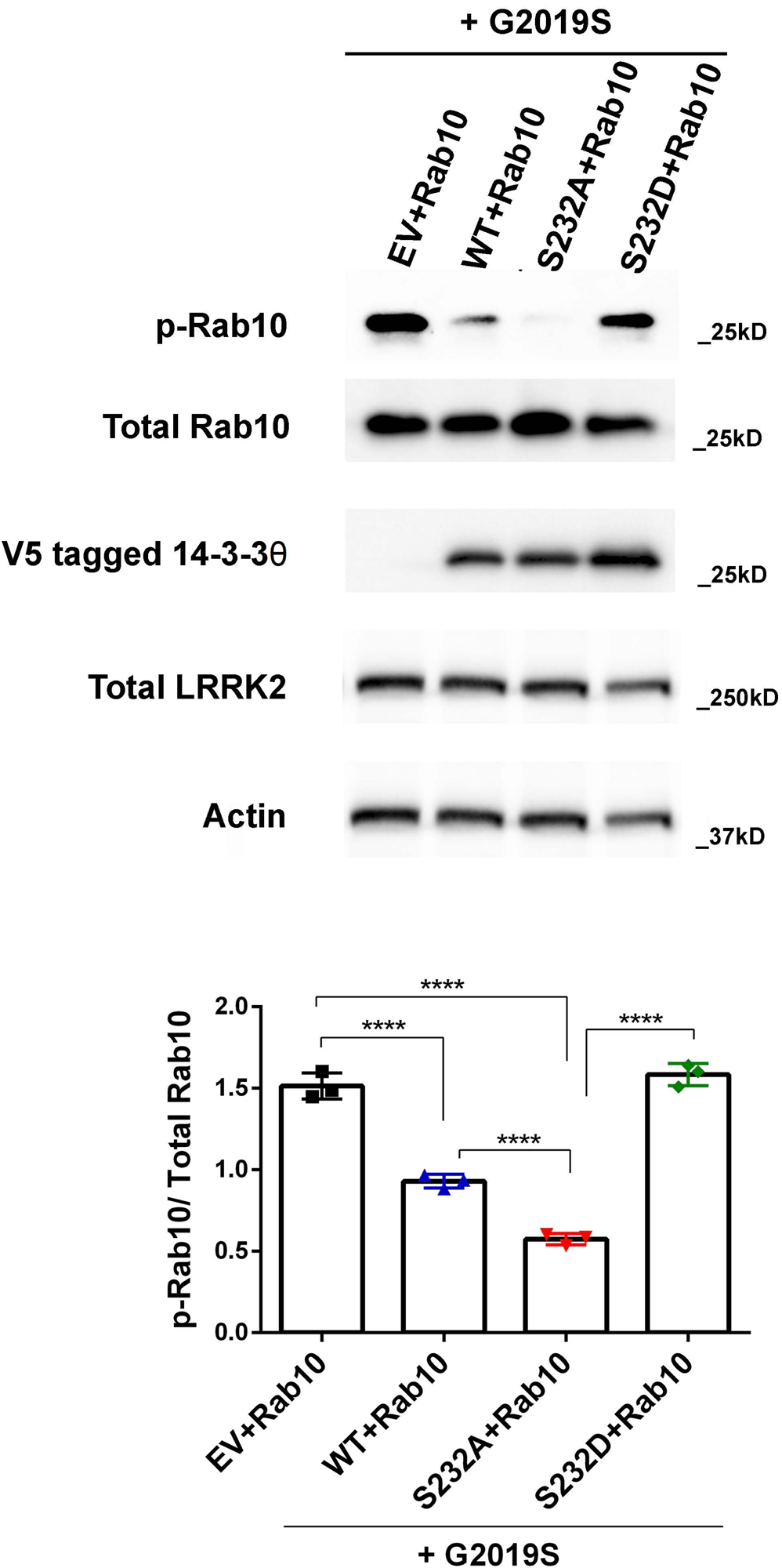
14-3-3θ phosphorylation reduces 14-3-3θ’s ability to reduce LRRK2-mediated Rab10 phosphorylation. Representative Western blot and quantification of p-T73 Rab10 levels. HA-tagged Rab10 and Flag-tagged G2019S were co-transfected with EV, V5-tagged WT, S232A, or S232D 14-3-3θ in HEK 293T cells. n = 3 independent rounds. ****p < 0.0001 (Tukey’s multiple comparison test)

### 14-3-3θ phosphorylation does not prevent binding to LRRK2

Our results above demonstrate that 14-3-3θ phosphorylation impacts the ability of 14-3-3θ to reduce the kinase activity of both WT and G2019S LRRK2 but not that of the R1441G mutant. We hypothesized that this differential effect could be related to differences in binding of 14-3-3θ mutants to different LRRK2 mutants. To test this hypothesis, we first measured the impact of the S232 mutants on S935 phosphorylation of WT, G2019S, and R1441G LRRK2. S910 and S935 have been shown to be key interaction sites between 14-3-3 proteins and LRRK2, and binding of 14-3-3s to LRRK2 prevents the dephosphorylation of these constitutively phosphorylated serines, S910 and S935 [12, 14]. As a result, phosphorylation at S935 is an indirect measure of 14-3-3/LRRK2 interactions. HEK293T cells were co-transfected with G2019S or R1441G LRRK2 along with WT or mutant 14-3-3θ. As expected, we found an increase in S935 phosphorylation in cells transfected with either LRRK2 mutant along with WT 14-3-3θ compared to control cells transfected with EV and LRRK2 mutant (Fig. 3A, B). S232A mutant similarly increased p-S935 LRRK2 signal in cells transfected with either G2019S or R1441G LRRK2 (Fig. 3A, B). Surprisingly, we found that the S232D mutant also promoted S935 phosphorylation of both G2019S and R1441G LRRK2 (Fig. 3A, B). Based on this finding, we concluded that 14-3-3θ phosphorylation did not impact the interaction between 14-3-3θ and LRRK2 mutants at least around the S935 site.

**Figure 3.**
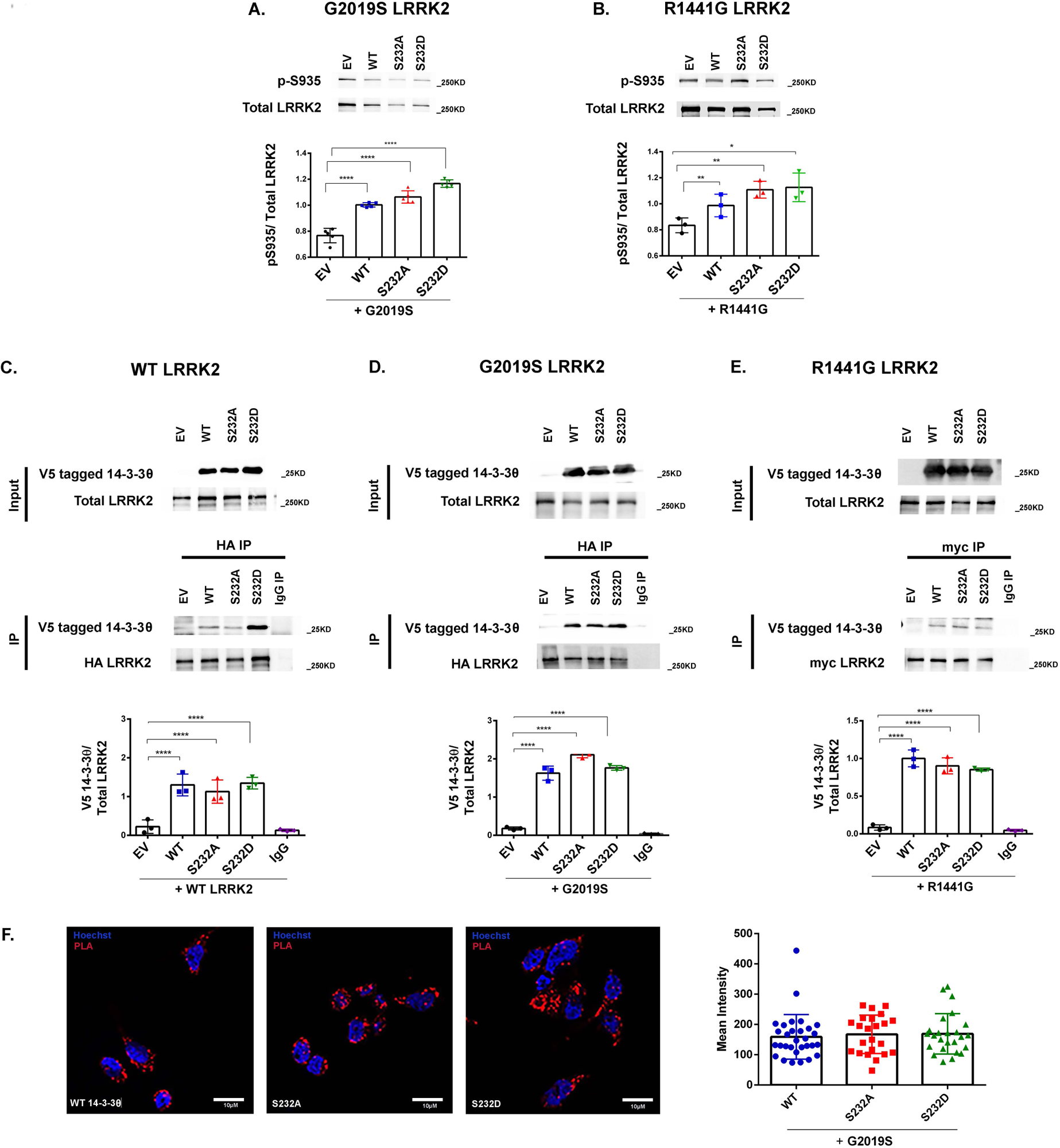
S232D phosphorylation does not cause dissociation of 14-3-3θ binding to LRRK2. **(A)** Representative Western blot and quantification of p-S935 LRRK2 levels in lysates of HEK 293T cells co-transfected with EV, WT, S232A, or S232D 14-3-3θ with G2019S LRRK2. n = 5 independent rounds. **** p < 0.0001 (Tukey’s multiple comparison test). **(B)** Representative Western blot and quantification of p-S935 LRRK2 levels in lysates of HEK 293T cells co-transfected with EV, WT, S232A, or S232D 14-3-3θ with R1441G LRRK2. n = 3 independent rounds. *p < 0.05, **p < 0.01 (Tukey’s multiple comparison test). **(C-E)** Cell lysates from HEK293T cells co-transfected with HA-tagged WT LRRK2 (C), HA-tagged G2019S LRRK2 (D), or c-myc-tagged R1441G LRRK2 (E) with EV, V5-tagged WT, S232A or S232D 14-3-3θ were immunoprecipitated with monoclonal antibodies against HA and c-myc, respectively. The resulting immunoprecipitants were analyzed for V5-tagged 14-3-3θ by Western blot. Lysates from cells transfected with S232A with HA-tagged WT LRRK2 (C), S232A with HA-tagged G2019S LRRK2 (D), or WT 14-3-3θ with c-myc-tagged R1441G LRRK2 (E) were used for IgG control. n = 3 independent rounds. ****p < 0.0001 (Tukey’s multiple comparison test). **F.** Representative PLA signal and quantification in HEK293T cells transfected with WT 14-3-3θ, S232A, or S232D along with G2019S LRRK2 Mean intensities for 30 cells per condition were analyzed with NIS elements analyzer software. n = 3 independent rounds; 10 cells/round.

We next performed co-immunoprecipitation assays to determine whether 14-3-3θ phosphorylation disrupted LRRK2 binding. We co-transfected epitope-tagged WT or mutant LRRK2 with V5-tagged WT or mutant 14-3-3θ into HEK 293T cells and then performed co-immunoprecipitation for LRRK2 and 14-3-3θ. In the case of WT LRRK2, we found that WT, S232A, and S232D 14-3-3θ showed similar levels of co-immunoprecipitation with WT LRRK2 (Fig. 3C). WT and mutant 14-3-3θ similarly were co-immunoprecipitated with G2019S LRRK2 and R1441G LRRK2 (Fig. 3D, E).

To verify our co-immunoprecipitation results, we next performed proximal ligation assay (PLA) between 14-3-3θ and LRRK2. HEK293T cells were transfected with G2019S LRRK2 along with WT or mutant 14-3-3θ, and then cells were fixed and underwent PLA using antibodies directed against LRRK2 and 14-3-3θ. As expected, WT 14-3-3θ showed binding to G2019S as determined by PLA (Fig. 3F). Cells transfected with either S232A or S232D 14-3-3θ mutants showed comparable PLA signal to those cells transfected with WT 14-3-3θ. Based on these assays, we concluded that 14-3-3θ phosphorylation did not cause a global dissociation between 14-3-3θ and either WT or mutant LRRK2.

### 14-3-3θ phosphorylation impacts the interaction of 14-3-3θ with LRRK2 at the C-terminal end

While our experiments demonstrated that 14-3-3θ phosphorylation did not cause a dissociation between 14-3-3θ and LRRK2, we next hypothesized that this phosphorylation could have caused more subtle changes in the 14-3-3θ/LRRK2 interaction that could have altered the conformational state of LRRK2 to affect its kinase activity. Besides S910, S935, and S1444, recent studies have shown that LRRK2 can interact with 14-3-3 proteins at its C-terminal end through residue T2524 [18]. This C-terminal end has been identified to fold back towards the kinase domain and thus regulate LRRK2 kinase activity, and phosphorylation at T2524 could potentially impact this conformational change [28]. Thus, we hypothesized that 14-3-3θ phosphorylation could impact its interaction with LRRK2 at this T2524 residue in order to impact kinase activity.

To test this hypothesis, we examined the impact of WT and mutant 14-3-3θ on LRRK2 kinase activity when T2524 was mutated to A. WT 14-3-3θ failed to reduce the kinase activity of the G2019S/T2524A LRRK2 double mutant as assessed by S1292 phosphorylation (Fig. 4A). Similarly, the S232A and S232D 14-3-3θ mutants failed to reduce S1292 phosphorylation of G2019S/T2524A LRRK2 (Fig. 4A). These findings were confirmed by testing for autophosphorylation at T1503 using an *in vitro* kinase assay after pulldown for LRRK2; neither WT nor mutant 14-3-3θ was able to reduce the kinase activity of G2019S/T2524A (Fig. 4B). No differences in co-immunoprecipitation were observed between WT, S232A, or S232D 14-3-3θ with the G2019S/T2524A double mutant (Fig. 4C). Based on these results, we conclude that the ability of WT and S232A 14-3-3θ to regulate LRRK2 kinase activity is dependent on 14-3-3θ’s interaction with LRRK2 via phosphorylation at the T2524 site.

**Figure 4.**
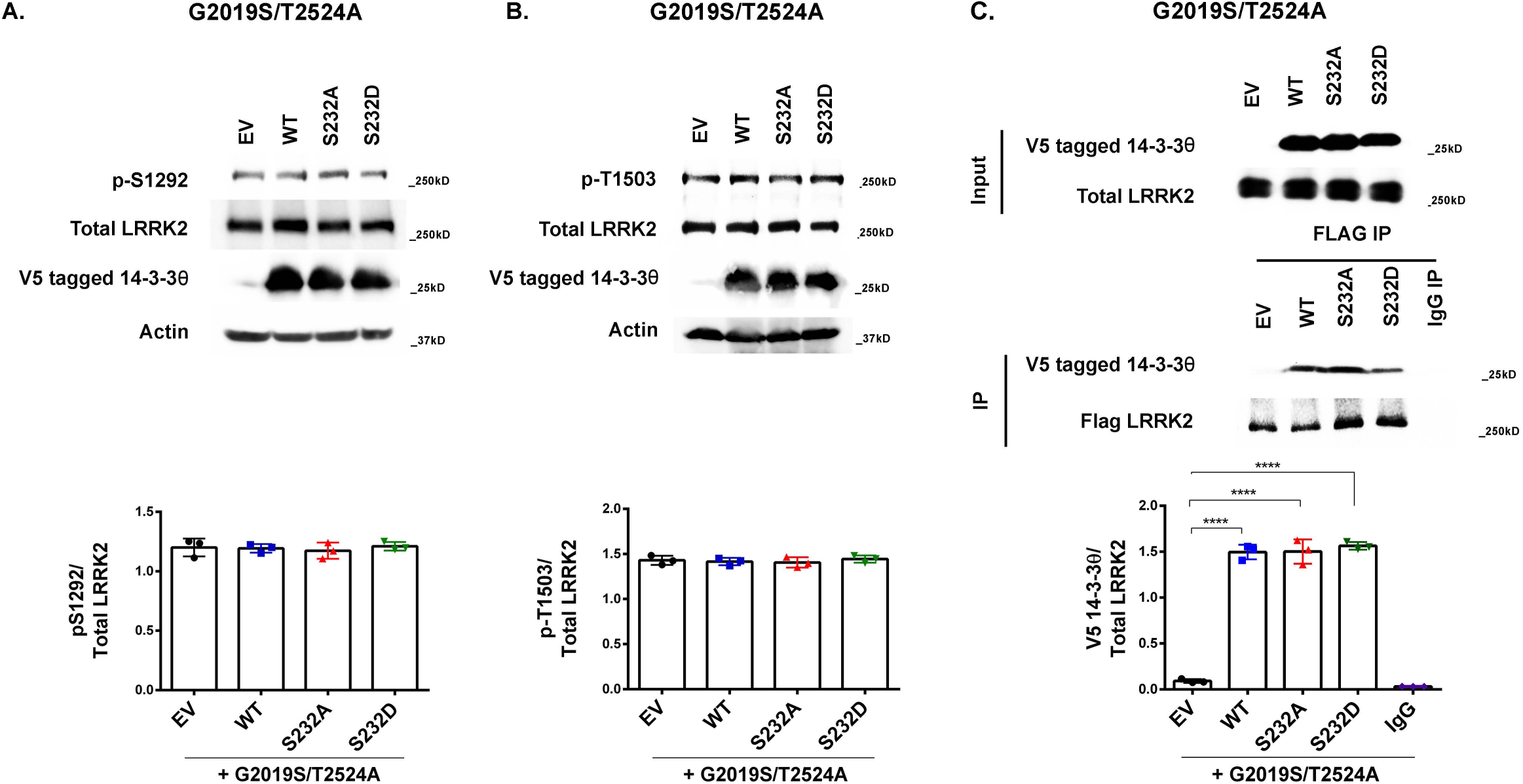
WT or mutant 14-3-3θ are unable to reduce autophosphorylation by G2019S/T2524A LRRK2 mutant. **(A)** Representative Western blot and quantification of p-S1292 LRRK2 levels in lysates of HEK 293T cells co-transfected with EV, V5 tagged WT, S232A, or S232D 14-3-3θ with the G2019S/T2524A double mutant. n = 3 independent rounds. **(B)** FLAG-tagged G2019S/T2524A LRRK2 was co-transfected with EV, WT, S232A or S232D 14-3-3θ into HEK 293T cells, and then LRRK2 was immunoprecipitated with FLAG antibody to isolate LRRK2 for kinase reactions. Following kinase reaction, immunoprecipitants were analyzed for p-T1503 and total LRRK2 by Western blot. n = 3 independent rounds. **(C)** Lysates from HEK293T cells co-transfected with FLAG-tagged G2019S/T2524A LRRK2 and either EV, V5-tagged WT, S232A, or S232D 14-3-3θ was immunoprecipitated FLAG antibody against LRRK2 and then probed for V5 to detect 14-3-3θ. Lysate from cells transfected with S232A with FLAG-tagged G2019S/T2524A LRRK2 was used for IgG control. ****p < 0.0001, n = 3 independent rounds.

### 14-3-3θ phosphorylation alters the binding pocket of 14-3-3θ

Based on these data, we hypothesized that phosphorylation at S232 likely causes conformational changes in the interaction between 14-3-3θ and LRRK2 at the C-terminal end so that 14-3-3θ is less effective at reducing LRRK2 kinase activity. To examine the impact of 14-3-3θ phosphorylation on the three dimensional structure of 14-3-3θ, we performed molecular dynamics simulations on both the unphosphorylated 14-3-3θ and 14-3-3θ phosphorylated at S232 (14-3-3θ^pS232^). As shown in Fig. 5A, snapshots of the trajectories reveal that the C-terminus of 14-3-3θ adopts a poorly ordered conformation. This observation is consistent with previous structural studies that show that the C-terminus of 14-3-3 proteins is unable to be resolved in crystal structures and is thus, inherently flexible and unstructured [29]. In contrast, phosphorylation of S232 induces a 90° kink in the C-terminus within 1 ns of the 14-3-3θ simulation (Fig. 5B). At 100 ns, the C-terminus adopts a stable conformation through contacts with the αE, αF, αG, and αH helices of 14-3-3θ.

**Figure 5.**
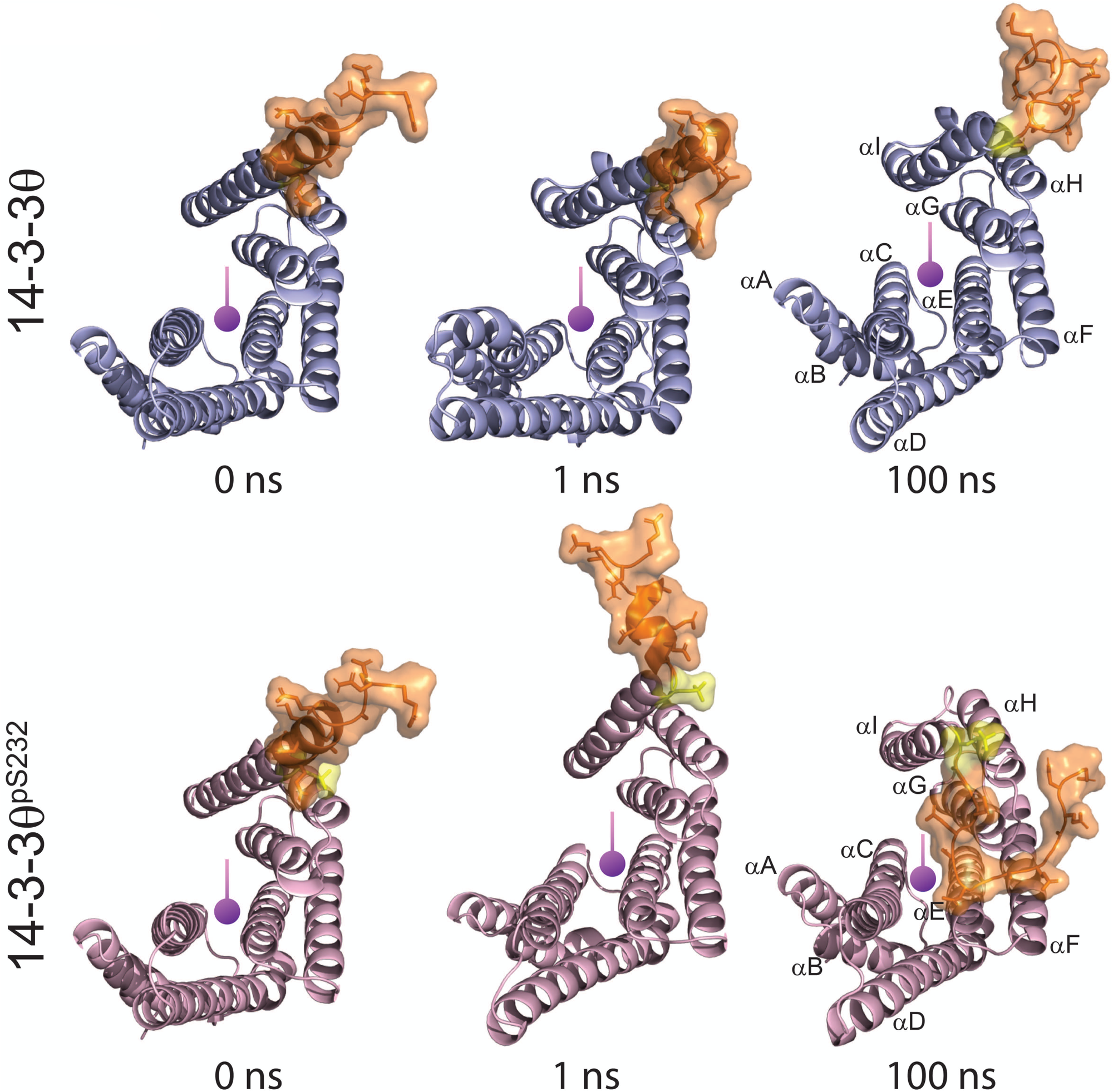
Molecular dynamics simulations reveal structural consequences of phosphorylation of 14-3-33θ at position S232. **(A, B)** Structures of 14-3-3θ (A) and 14-3-3θ^p232^ (B) were energetically minimized and subjected to molecular dynamics simulations of 100 ns. Snapshots of the trajectories at 0 ns, 1ns, and 100ns are shown with the canonical binding pocket of 14-3-3θ indicated by a purple indicator with each α-helix labeled.

The structural rearrangement of the C-terminus caused by phosphorylation of S232 has a dramatic effect on local electronic and steric environment of the canonical 14-3-3θ binding pocket. As shown in Fig. 6A and 6B, the structural rearrangement and packing of the C-terminus of 14-3-3θ^pS232^ alters the electrostatic surface, especially at its interface with αE, αF, αG, and αH. The findings that the LRRK2/14-3-3θ interaction is facilitied by the C-terminus of LRRK2 via residue T2524 suggest that the interaction between LRRK2 and 14-3-3θ is of the mode III variety. Mode III binding is when 14-3-3 binds a phosphorylated residue in the unstructured C-terminus of a target protein [30], in this case pT2524 of LRRK2. Based on previous studies analyzing binding mode III of 14-3-3 proteins [30, 31], we are able to predict the orientation of the unstructured C-terminus of LRRK2 in the 14-3-3θ binding pocket. The relative positions of the 14-3-3θ side chains that interact with LRRK2 amino acid residues just downstream of pT2524 (i.e. towards the C-terminus end) remain virtually unchanged upon phosphorylation of S232 (Fig. 6C). In contrast, the 14-3-3θ side chains that would interact with LRRK2 residues just upstream (i.e. towards the N-terminal direction) of pT2524 (Fig. 6D) show significant rearrangement within the binding pocket. Interestingly, several side chains known to play critical roles in the interactions between 14-3-3s and other proteins, irrespective of binding mode, adopt new ionic interactions upon S232 phosphorylation throughout the 14-3-3θ binding pocket (Fig. 6E, F). These residues include but are not limited to R49, R56, and R127. Taken together, the structural rearrangement that we observed upon phosphorylation of S232 could offer a mechanistic explanation of our biological data.

**Figure 6.**
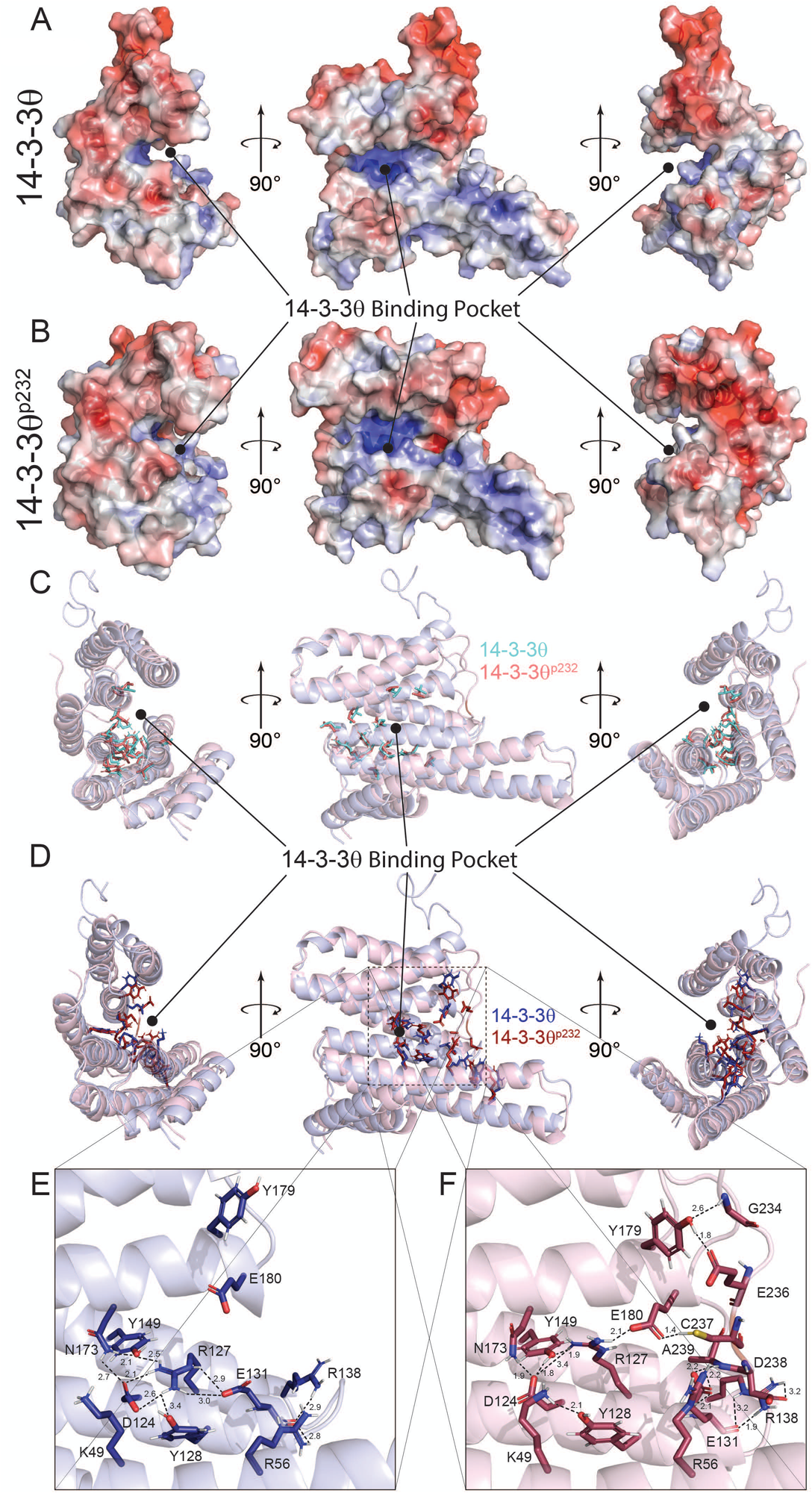
Phosphorylation of 14-3-3θ at position S232 causes a partial rearrangement of its canonical binding pocket. **(A, B)** Electrostatic surfaces of 14-3-3θ (A) and 14-3-3θ^p232^ (B) shown for the 100 ns snapshot of the molecular dynamics simulation with blue indicating positive surfaces charge and red indicating negative surface charges. **(C, D)** Superposition of 14-3-3θ and 14-3-3θ^p232^ with the side chains comprising the 14-3-3θ that maintain their relative position (C) and undergo a dramatic rearrangement (D) after phosphorylation. **(E, F)** Panels are magnified images of the binding pocket rearrangement without phosphorylation (E) and with phosphorylation (F). Dotted lines indicate ionic interaction and the presence of hydrogen bonds with the distance between the two atoms indicated in angstroms. The canonical binding pocket of 14-3-3θ is labeled.

## DISCUSSION

14-3-3 proteins, including the 14-3-3θ isoform, are major regulators for LRRK2 kinase activity [17, 19]. We previously observed a significant increase in 14-3-3θ phosphorylation in detergent-insoluble brain fractions from human PD subjects [23]. In this study, we examined the potential impact of 14-3-3θ phosphorylation on its ability to regulate the kinase activity of different LRRK2 mutants. We found that WT 14-3-3θ and the non-phosphorylatable S232A mutant reduced the kinase activity of both WT and G2019S LRRK2, while the phosphomimetic S232D mutant failed to reduce the kinase activity of either WT or G2019S LRRK2, as determined through LRRK2 autophosphorylation and Rab10 phosphorylation. However, S232 phosphorylation did not cause a global dissociation of 14-3-3θ binding to LRRK2, as determined by S935 phosphorylation, co-immunoprecipitation, or PLA studies. Instead, our findings showed that S232 phosphorylation caused structural changes in the binding pocket of 14-3-3θ that affected 14-3-3θ’s interaction with LRRK2 involving T2524 at the C-terminal end. Our results suggest that this altered interaction with LRRK2 at its C-terminal end impacted the ability of 14-3-3θ to regulate kinase activity.

Several different 14-3-3 interaction sites have been described for LRRK2, including S910, S935, S1444 [12, 14, 19], and most recently T2524 in the C-terminal end [18], and interactions of these sites with 14-3-3s is regulated by phosphorylation at these sites [12, 14, 18, 19]. Several studies have demonstrated the significance of the C-terminal end in regulating the kinase domain of LRRK2 [28, 32]. Various deletions of the C-terminal region abolish the kinase activity of LRRK2 and can also disrupt microtubule association and 14-3-3 interaction [32, 33]. Structural studies have suggested that the C-terminal end forms a helical structure that can interact with the kinase domain to tether the kinase domain into an active conformation [28]. Phosphorylation of T2524 is predicted to regulate this interaction between the C-terminal end and the kinase domain and thus impact kinase activity [18, 28]. Binding of 14-3-3s to T2524 is dependent on phosphorylation at T2524 [18],and this phosphorylation site at the C-terminal end of LRRK2 is consistent with one of the canonical 14-3-3 binding motifs [18]. Based on our findings here, we predict that upon T2524 phosphorylation, 14-3-3θ interacts with LRRK2 at this C-terminal site to promote conformational changes in the interaction between the C-terminus and the kinase domain and thus reduce LRRK2 kinase activity. When we introduced the T2524A mutation in G2019S, we found that WT and S232A 14-3-3θ were no longer capable of reducing the kinase activity of this double mutant LRRK2. Therefore, we conclude that the interaction between 14-3-3θ and LRRK2 at the phosphorylated T2524 residue is important for 14-3-3θ to regulate LRRK2 kinase activity. 14-3-3θ phosphorylation at S232 likely prevents the normal interaction between 14-3-3θ and LRRK2 at T2524 so that 14-3-3θ is less effective at reducing LRRK2 kinase activity. In PD in which excess 14-3-3θ phosphorylation has been observed in the brain [23], aberrant 14-3-3θ phosphorylation at S232 could thus promote LRRK2 kinase activity and toxicity.

Molecular modeling revealed that 14-3-3θ phosphorylation at S232 causes structural rearrangements of the canonical binding pocket. Assuming a mode III type of interaction between 14-3-3θ and the C-terminus of LRRK2, this structural change in the binding pocket would explain our biological observations. While we observe steric and ionic rearrangements of side chains in the canonical binding pocket upon phosphorylation of 14-3-3θ at S232, there are aspects of the binding pocket that remain unaltered. We posit that this would lead not to a complete inhibition of the interaction between 14-3-3θ and the C-terminus of LRRK2, but a modulation of the binding properties between the two proteins.

Phosphorylation of 14-3-3 proteins is a key mechanism for the regulation of 14-3-3 function [34, 35]. Three separate conserved 14-3-3 phosphorylation sites have been described, including S58/59, S184/185, and S232/T233, with 14-3-3 isoforms differing in which phosphorylation sites are present [24, 36–38]. Here we shown that phosphorylation at S232 is critical to the ability of 14-3-3θ to regulate LRRK2 interaction and kinase activity. Civiero *et al.* 2017 have identified that 14-3-3 phosphorylation at S59 can also regulate its impact of LRRK2 function [39]. PAK6 can phosphorylate the 14-3-3γ isoform, which then disrupts the interaction of 14-3-3γ with LRRK2 and reduces LRRK2 phosphorylation at S935 [39]. Alterations in 14-3-3 phosphorylation at several sites are likely critical mechanisms for regulating the interaction and function of LRRK2.

Mechanisms underlying 14-3-3θ phosphorylation at S232 are not well understood. Our work has shown that both alpha-synuclein overexpression and the mitochondrial toxin rotenone, which is associated with increased PD risk [40–42], can induce S232 phosphorylation *in vitro* [24]. These and other triggers could promote excessive LRRK2 kinase activity via 14-3-3θ phosphorylation in order to promote neurodegeneration in PD. Indeed, rotenone has been shown to disrupt the interaction of 14-3-3s with LRRK2 in rodent models and promote LRRK2 kinase activity [43].

We observed differential effects of 14-3-3θ phosphorylation on different LRRK2 mutants. S232 phosphorylation showed clear impact on the ability of 14-3-3θ to regulate both WT and G2019S LRRK2 kinase activity; however, we did not observe any differences between WT, S232A, and S232D 14-3-3θ in their ability to reduce the kinase activity of the R1441G LRRK2 mutant. All three 14-3-3θ constructs were capable of reducing R1441G LRRK2 kinase activity in our assays. G2019S is within the activation loop of the kinase domain of LRRK2 and is therefore in a position to be regulated by the C-terminal helix/14-3-3θ interaction. In contrast, R1441 is in the ROC domain, and mutations in this site reduce GTPase activity and have less pronounced effects on kinase activity [44]. Our findings suggest that S232 phosphorylation has no impact on the ability of 14-3-3θ to interact with the R1441G mutant and impact kinase activity. Interestingly, the R1441G mutation is near another critical 14-3-3 binding site, S1444, which is dependent on phosphorylation by PKA; access to this phosphorylation site by PKA is reduced by mutations at the R1441 site and reduce the affinity for 14-3-3s [19]. We predict that S232 phosphorylation will not affect all LRRK2 mutants similarly but will impact those mutations within kinase domain more significantly.

In conclusion, 14-3-3θ phosphorylation at S232 disrupts the ability of 14-3-3θ to reduce the kinase activity of WT and G2019S LRRK2. This effect is likely mediated by 14-3-3θ phosphorylation altering the binding interaction for the C-terminal end of LRRK2, which can fold back to regulate the kinase domain of LRRK2. Increased 14-3-3θ phosphorylation observed in human PD could thus promote LRRK2 toxicity by dysregulation of LRRK2 kinase activity.

## METHODS

### Transfection

HEK293T cells were grown in DMEM media supplemented with 10% fetal bovine serum and 1% penicillin/streptomycin. When the cells were 70-80% confluent, they were transfected with WT or mutant LRRK2, WT or mutant 14-3-3θ, and/or Rab10 using Superfect or Polyfect transfection reagent (Qiagen, Germantown, MD) following the manufacturer’s protocol. After transfection, the cells were incubated in fresh media for 48 hours before harvesting the protein.

### Immunoblotting

Cell lysates were washed with PBS and then harvested using RIPA lysis buffer (Thermo Fisher Scientific, Waltham, MA) supplemented with protease inhibitor and phosphatase inhibitor (Pierce, Rockford, IL). The cells were kept on ice for 15 minutes and then sonicated for 10 seconds on ice. The lysates were centrifuged at 15000g for 10 minutes, and the supernatant was collected. Protein concentrations were measured by BCA assay kit (Pierce).

For Western blots to detect 14-3-3θ or Rab10, 50 µg of protein per sample was boiled in DTT sample loading buffer (0.25 M Tris–HCl, pH 6.8, 8% SDS, 200 mM DTT, 30% glycerol, and bromophenol blue) and resolved on 12% SDS polyacrylamide gel. Proteins were transferred to nitrocellulose membranes at 100V for 1 hour. For Western blots to detect p-S1292, p-T1503, or total LRRK2, 50 µg of protein was loaded per sample to 7.5% gel and transferred overnight to nitrocellulose membrane at 30V for 16 hours at 4°C. Membranes were then blocked in 5% non-fat milk solution prior to the incubation in primary antibodies (Table1) overnight at 4°C. Primary antibodies against p-LRRK2 and total LRRK2 were diluted in EMD signal boost solution 1 (Millipore Sigma, Burlington, MA). After washing, all membranes were incubated with HRP-conjugated goat anti-mouse or anti-rabbit secondary antibody (Jackson ImmunoResearch, West Grove, PA; 1:2000 dilution) corresponding to the primary antibody and developed with the enhanced chemiluminescence method. Images were scanned using the Bio-Rad Chemidoc Imaging System.

For detection of p-S935 LRRK2, PVDF membrane was used during transfer, and the membrane was incubated in Odyssey blocking buffer (LI-COR Biosciences, Lincoln, NE). The primary antibodies (Table 1) were diluted in 1:1 Odyssey buffer and EMD signal boost solution 1. After washing, IRDye 800CW Donkey anti Rabbit IgG (LI-COR Biosciences # 926-32212) and IRDye 680LT Donkey polyclonal anti-mouse IgG (LI-COR #926-68023) secondary antibodies were incubated with the membrane. The blot for p-S935 was developed using LI-COR Odessey CLx imager and analyzed with Image studio version 5.2 software.

### Immunoprecipitation

HA-tagged WT LRRK2 or G2019S or c-myc-tagged R1441G or FLAG-tagged T2524A-G2019S along with WT or mutant 14-3-3θ were co-transfected in HEK 293T cells prior to co-immunoprecipitation assay. 25 µl slurry of Protein G Dynabeads (Invitrogen, Carlsbad, CA) was incubated overnight with 4 µg mouse HA antibody or c-myc antibody (Table 1) respectively, at 4°C. 500 µg of cell lysate per sample was incubated with antibody-conjugated beads for 30 minutes at room temperature. The protein/bead complex were washed five times in PBS with 0.02% Tween, boiled in DTT sample loading buffer, and then divided in two equal portions. One half of sample was run for total LRRK2 using a 7.5% gel, and the other half was run for 14-3-3θ using a 12% gel. After transfer, membranes were probed for LRRK2 and V5-tagged 14-3-3θ.

### LRRK2 kinase assay

Epitope-tagged WT, G2019S, R1441G, or G2019S/T2524A-G2019S with or without V5-tagged 14-3-3θ mutants were transfected in HEK293T cells. After 48 hours, cell lysates were prepared and then incubated with respective epitope tag antibody (Table 1) that was conjugated to protein G Dynabeads for 30 minutes. Beads were washed twice with PBS with 500 mM NaCl and then resuspended in kinase buffer (10 mM Tris pH 7.4, 0.1 mM EGTA, 20 mM MgCl2, 0.1 mM ATP). The samples were then shaken at 30°C for 30 minutes. The kinase reaction was terminated by incubating samples on ice, and beads were then incubated in Laemmli buffer at 95°C for 10 minutes. Sample was then run on a 7.5% SDS-acrylamide gel, transferred to nitrocellulose, and probed for phosphorylated LRRK2 at T1503 using a primary rabbit antibody against phospho-T1503 and for total LRRK2 (Table 1).

### Proximal Ligation assay

HEK293T cells were transfected with G2019S LRRK2 and WT or mutant 14-3-3θ using Polyfect reagent. 48 hours after transfection, cells were fixed and incubated with primary antibodies against 14-3-3θ and against LRRK2 (UDD3, Abcam) overnight on shaker at 4°C. PLUS and MINUS PLA probes were diluted in the Duolink Antibody diluent in 1:5 ratio with blocking buffer from the PLA kit (Millipore Sigma). The coverslips were then incubated in PLA probe solution for 1 hour at 37°C in humidity chamber. After washing with buffer A at RT, the coverslips were incubated in Duolink ligation buffer with ligase enzyme for 45 minutes at 37°C in humidity chamber. After washing, the cells on coverslips were incubated in the amplification buffer for 100 minutes at 37°C in humidity chamber. The coverslips were washed with wash buffer B and mounted in Duolink Mounting Media with DAPI. Red florescence detecting interaction was detected using Nikon Eclipse Ti2 scanning confocal microscope. The images were analyzed for mean intensity per cell using NIS-Elements analysis software. Mean intensity of randomly selected thirty cells per condition were used for the statistical analysis.

### Molecular modeling

The three-dimensional structure of 14-3-3θ used for the molecular dynamics simulations was generated by using AlphaFold [45] to predict the structural elements missing from the previously solved crystal structure (PBD: 5IQP) [46]. Phosphorylation of residue S232 was modeled in Pymol [47] using the PyTMs plugin [48]. All molecular dynamics simulations were performed using GROMACS 2019 software with the GROMOS 54A7 force field [49] and SPC216 water model. Specifically, each protein was solvated with 36,080 SPC216 explicit water molecules and placed in the center of a cubic box of 104 × 104 × 104 Å^3^. Each system was neutralized by adding the appropriate number of Na^+^ ions. Energy minimization of each structure was followed by a two-step equilibration (i.e. NVT equilibration followed by NPT equilibration). The temperature of each system was controlled through velocity rescaling[50] at 300 K with a time constant of 0.1 picosecond. The pressure of each system was controlled using the Parrinello-Rahman barostat [51] and set to 1 bar. The particle mesh Ewald algorithm [52] was used to calculate long-range electrostatics while a cutoff of 1.0 nM was used for short-range electrostatics and van der Waals’ interactions. Molecular dynamics simulations of 100 ns were performed for 14-3-3θ and 14-3-3θ^pS232^ with linear constraint solver (LINCS) [53] constraints for all bonds. Frames were recorded every 2 ps. The backbone RMSD was monitored over the production run of each protein to ensure the stability and convergence of the simulated trajectories. Analysis of the electrostatic surfaces were performed using the APBS (Adaptive Poisson-Boltzmann Solver) software [54].

### Statistical analysis

Graph Pad Prism 9.4 (La Jolla, CA) has been for statistical analysis of experiments. Experiments were analyzed by 1-way ANOVA, followed by post-hoc Dunnett’s or Tukey’s multiple comparison test.

## Acknowledgements

This study was supported by NIH R01NS112203.

## References

1. Healy, D.G., et al., Phenotype, genotype, and worldwide genetic penetrance of LRRK2-associated Parkinson’s disease: a case-control study. Lancet Neurol, 2008. 7(7): p. 583–90.

2. Paisan-Ruiz, C., et al., Cloning of the gene containing mutations that cause PARK8-linked Parkinson’s disease. Neuron, 2004. 44(4): p. 595–600.

3. Zimprich, A., et al., Mutations in LRRK2 cause autosomal-dominant parkinsonism with pleomorphic pathology. Neuron, 2004. 44(4): p. 601–7.

4. Myasnikov, A., et al., Structural analysis of the full-length human LRRK2. Cell, 2021. 184(13): p. 3519–3527 e10.

5. Price, A., et al., The LRRK2 signalling system. Cell Tissue Res, 2018. 373(1): p. 39–50.

6. Greggio, E., et al., Kinase activity is required for the toxic effects of mutant LRRK2/dardarin. Neurobiol Dis, 2006. 23(2): p. 329–41.

7. Smith, W.W., et al., Kinase activity of mutant LRRK2 mediates neuronal toxicity. Nat Neurosci, 2006. 9(10): p. 1231–3.

8. West, A.B., et al., Parkinson’s disease-associated mutations in LRRK2 link enhanced GTP-binding and kinase activities to neuronal toxicity. Hum Mol Genet, 2007. 16(2): p. 223–32.

9. Wang, X., et al., Understanding LRRK2 kinase activity in preclinical models and human subjects through quantitative analysis of LRRK2 and pT73 Rab10. Sci Rep, 2021. 11(1): p. 12900.

10. Correia Guedes, L., et al., Worldwide frequency of G2019S LRRK2 mutation in Parkinson’s disease: a systematic review. Parkinsonism Relat Disord, 2010. 16(4): p. 237–42.

11. Williams-Gray, C.H., et al., Prevalence of the LRRK2 G2019S mutation in a UK community based idiopathic Parkinson’s disease cohort. J Neurol Neurosurg Psychiatry, 2006. 77(5): p. 665–7.

12. Li, X., et al., Phosphorylation-dependent 14-3-3 binding to LRRK2 is impaired by common mutations of familial Parkinson’s disease. PLoS One, 2011. 6(3): p. e17153.

13. Dzamko, N., et al., Inhibition of LRRK2 kinase activity leads to dephosphorylation of Ser(910)/Ser(935), disruption of 14-3-3 binding and altered cytoplasmic localization. Biochem J, 2010. 430(3): p. 405–13.

14. Nichols, R.J., et al., 14-3-3 binding to LRRK2 is disrupted by multiple Parkinson’s disease-associated mutations and regulates cytoplasmic localization. Biochem J, 2010. 430(3): p. 393–404.

15. Pair, F.S. and T.A. Yacoubian, 14-3-3 Proteins: Novel Pharmacological Targets in Neurodegenerative Diseases. Trends Pharmacol Sci, 2021. 42(4): p. 226–238.

16. Giusto, E., et al., Pathways to Parkinson’s disease: a spotlight on 14-3-3 proteins. npj Parkinson’s Disease, 2021. 7(1): p. 85.

17. Lavalley, N.J., et al., 14-3-3 Proteins regulate mutant LRRK2 kinase activity and neurite shortening. Hum Mol Genet, 2016. 25(1): p. 109–22.

18. Manschwetus, J.T., et al., Binding of the Human 14-3-3 Isoforms to Distinct Sites in the Leucine-Rich Repeat Kinase 2. Front Neurosci, 2020. 14: p. 302.

19. Muda, K., et al., Parkinson-related LRRK2 mutation R1441C/G/H impairs PKA phosphorylation of LRRK2 and disrupts its interaction with 14-3-3. Proc Natl Acad Sci U S A, 2014. 111(1): p. E34–43.

20. Kawamoto, Y., et al., 14-3-3 proteins in Lewy bodies in Parkinson disease and diffuse Lewy body disease brains. J Neuropathol Exp Neurol, 2002. 61(3): p. 245–53.

21. Wakabayashi, K., et al., 14-3-3 protein sigma isoform co-localizes with phosphorylated α-synuclein in Lewy bodies and Lewy neurites in patients with Lewy body disease. Neuroscience Letters, 2018. 674: p. 171–175.

22. Berg, D., O. Riess, and A. Bornemann, Specification of 14-3-3 proteins in Lewy bodies. Annals of Neurology, 2003. 54(1): p. 135–135.

23. McFerrin, M.B., et al., Dysregulation of 14-3-3 proteins in neurodegenerative diseases with Lewy body or Alzheimer pathology. Ann Clin Transl Neurol, 2017. 4(7): p. 466–477.

24. Slone, S.R., et al., Increased 14-3-3 phosphorylation observed in Parkinson’s disease reduces neuroprotective potential of 14-3-3 proteins. Neurobiol Dis, 2015. 79: p. 1–13.

25. Liu, Z., et al., LRRK2 and Rab10 coordinate macropinocytosis to mediate immunological responses in phagocytes. Embo j, 2020. 39(20): p. e104862.

26. Malik, A.U., et al., Deciphering the LRRK code: LRRK1 and LRRK2 phosphorylate distinct Rab proteins and are regulated by diverse mechanisms. Biochem J, 2021. 478(3): p. 553–578.

27. Steger, M., et al., Phosphoproteomics reveals that Parkinson’s disease kinase LRRK2 regulates a subset of Rab GTPases. Elife, 2016. 5.

28. Deniston, C.K., et al., Structure of LRRK2 in Parkinson’s disease and model for microtubule interaction. Nature, 2020. 588(7837): p. 344-349.

29. Yang, X., et al., Structural basis for protein-protein interactions in the 14-3-3 protein family. Proc Natl Acad Sci U S A, 2006. 103(46): p. 17237–42.

30. Ganguly, S., et al., Melatonin synthesis: 14-3-3-dependent activation and inhibition of arylalkylamine N-acetyltransferase mediated by phosphoserine-205. Proc Natl Acad Sci U S A, 2005. 102(4): p. 1222–7.

31. Coblitz, B., et al., C-terminal binding: an expanded repertoire and function of 14-3-3 proteins. FEBS Lett, 2006. 580(6): p. 1531–5.

32. Rudenko, I.N., et al., The G2385R variant of leucine-rich repeat kinase 2 associated with Parkinson’s disease is a partial loss-of-function mutation. Biochem J, 2012. 446(1): p. 99–111.

33. Cresto, N., et al., The C-terminal domain of LRRK2 with the G2019S mutation is sufficient to produce neurodegeneration of dopaminergic neurons in vivo. Neurobiol Dis, 2020. 134: p. 104614.

34. Aitken, A., Functional specificity in 14-3-3 isoform interactions through dimer formation and phosphorylation. Chromosome location of mammalian isoforms and variants. Plant Mol Biol, 2002. 50(6): p. 993–1010.

35. Aitken, A., 14-3-3 proteins: a historic overview. Semin Cancer Biol, 2006. 16(3): p. 162–72.

36. Obsilova, V., et al., 14-3-3zeta C-terminal stretch changes its conformation upon ligand binding and phosphorylation at Thr232. J Biol Chem, 2004. 279(6): p. 4531–40.

37. Sunayama, J., et al., JNK antagonizes Akt-mediated survival signals by phosphorylating 14-3-3. J Cell Biol, 2005. 170(2): p. 295–304.

38. Woodcock, J.M., et al., The dimeric versus monomeric status of 14-3-3zeta is controlled by phosphorylation of Ser58 at the dimer interface. J Biol Chem, 2003. 278(38): p. 36323–7.

39. Civiero, L., et al., PAK6 Phosphorylates 14-3-3γ to Regulate Steady State Phosphorylation of LRRK2. Front Mol Neurosci, 2017. 10: p. 417.

40. Liu, B., H.M. Gao, and J.S. Hong, Parkinson’s disease and exposure to infectious agents and pesticides and the occurrence of brain injuries: role of neuroinflammation. Environ Health Perspect, 2003. 111(8): p. 1065–73.

41. Tanner, C.M., et al., Rotenone, paraquat, and Parkinson’s disease. Environ Health Perspect, 2011. 119(6): p. 866–72.

42. Sherer, T.B., et al., Mechanism of toxicity in rotenone models of Parkinson’s disease. J Neurosci, 2003. 23(34): p. 10756–64.

43. Di Maio, R., et al., LRRK2 activation in idiopathic Parkinson’s disease. Sci Transl Med, 2018. 10(451).

44. Lewis, P.A., et al., The R1441C mutation of LRRK2 disrupts GTP hydrolysis. Biochem Biophys Res Commun, 2007. 357(3): p. 668–71.

45. Jumper, J., et al., Highly accurate protein structure prediction with AlphaFold. Nature, 2021. 596(7873): p. 583–589.

46. Xiao, B., et al., Structure of a 14-3-3 protein and implications for coordination of multiple signalling pathways. Nature, 1995. 376(6536): p. 188–91.

47. Delano, W.L., The PyMOL Molecular Graphics System, Version 2.0 Schrödinger, LLC.

48. Warnecke, A., et al., PyTMs: a useful PyMOL plugin for modeling common post-translational modifications. BMC Bioinformatics, 2014. 15(1): p. 370.

49. Huang, W., Z. Lin, and W.F. van Gunsteren, Validation of the GROMOS 54A7 Force Field with Respect to beta-Peptide Folding. J Chem Theory Comput, 2011. 7(5): p. 1237–43.

50. Bussi, G., D. Donadio, and M. Parrinello, Canonical sampling through velocity rescaling. Journal of Chemical Physics, 2007. 126(1).

51. Parrinello, M. and A. Rahman, Polymorphic transitions in single crystals: A new molecular dynamics method. Journal of Applied Physics 1981. 52(12): p. 7182–7190.

52. Essmann, U., et al., A Smooth Particle Mesh Ewald Method. Journal of Chemical Physics, 1995. 103(19): p. 8577–8593.

53. Hess, B., et al., LINCS: A linear constraint solver for molecular simulations. Journal of Computational Chemistry, 1997. 18(12): p. 1463–1472.

54. Jurrus, E., et al., Improvements to the APBS biomolecular solvation software suite. Protein Sci, 2018. 27(1): p. 112–128.

